# To FID or not to FID: Applying GANs for MRI Image Generation in HPC

**DOI:** 10.1101/2024.09.27.615343

**Authors:** Beatriz Cepa, Cláudia Brito, António Sousa

## Abstract

With the rapid growth of Deep Learning models and neural networks, the medical data available for training – which is already significantly less than other types of data – is becoming scarce. For that purpose, Generative Adversarial Networks (GANs) have received increased attention due to their ability to synthesize new realistic images. Our preliminary work shows promising results for brain MRI images; however, there is a need to distribute the workload, which can be supported by High-Performance Computing (HPC) environments. In this paper, we generate 256*×*256 MRI images of the brain in a distributed setting. We obtained an FID_RadImageNet_ of 10.67 for the DCGAN and 23.54 for the WGAN-GP, which are consistent with results reported in several works published in this scope. This allows us to conclude that distributing the GAN generation process is a viable option to overcome the computational constraints imposed by these models and, therefore, facilitate the generation of new data for training purposes.

## I. Introduction

Medical image analysis aims to acquire information about the medical condition of a patient in a non-invasive way [1]. The use of Machine/Deep Learning (ML/DL) models to automatize this task has become increasingly popular since analyzing images manually requires an astounding effort from medical professionals and is very time-consuming [2]. For these models to perform well, they need large amounts of training data, which, in the medical domain, is often difficult to obtain due to privacy issues and time-consuming annotations [2], [3]. In addition, although there are several public medical datasets available for research use, they are smaller than other non-medical datasets [4] – for instance, while *ImageNet* [5] contains more than 14 million images, *RadImageNet* [6] is composed of 5 million images. Thus, the need has emerged to explore new methods of obtaining more data.

For that purpose, Generative Adversarial Networks (GANs) have received increased attention in tasks such as segmentation and classification due to their ability to synthesize new realistic images [1], [4]. Our preliminary work corroborates this claim by showing promising results for brain MRI images; however, we found the model computationally heavy, resulting in high CPU running times [7]. This is a result of the GAN learning process, which might be compared to a two-player game between two models: the Generator, trying to generate new images resembling real ones, and the Discriminator, which discriminates between real and synthetic samples [8]. One key takeaway from the preliminary work was that resorting to High-Performance Computing (HPC) environments would support the need for workload distribution and benefit the model training time and stability [7].

Furthermore, a second takeaway from previous work is that there is no perfect metric to evaluate the output of these generative models (*i.e*., a performance metric besides human evaluation should be used for scientific evaluations) [8]–[10]. Although works have been proposed showing that Fréchet Inception Distance (FID) can be a possible metric to use, it does not account for the fidelity and diversity of images in a set. With this, better-resolution images may have a lower FID even if the diversity of images in the set is lower than in a set with more diverse images and lower resolution [11]. Moreover, FID is computed with InceptionV3 [12] trained with the ImageNet dataset [5], in which the images do not translate into medical images [1], [3], [4], [9].

Backed by these insights, we re-implement the model used in our previous work (*i.e*., *DCGAN*), and also re-implement a *WGAN-GP* [13] to generate MRI images of the brain in a larger distributed setting. Both models were implemented in TensorFlow and resorted to the `MultiWorkerMirroredStrategy` module, which splits the model between the several GPU-enabled nodes.

A thorough evaluation was conducted by leveraging the two models and the 2020 and 2021 BraTS datasets [14]–[17] in an HPC infrastructure. Distinct from other works [1]– [4], [18], our models generate 256*×*256 images and have reached the lowest FID value of approximately 10.67 with the RadImageNet training weights.

## II. Background

### A. DCGAN

Deep Convolutional GANs (DCGANs) were introduced by Radford et al. [19] and quickly became one of the most used GANs in medical image analysis [4] due to their ability to generate higher-quality images. They rely on fractional-strided convolutions (also named transposed convolutions) on the generator and strided convolutions on the discriminator to generate 64*×*64 realistic images. In our previous work [7], we implemented this architecture with the Chainer framework [20] and obtained promising results in generating 256*×*256 MRI images of the brain in a single-node CPU.

### B. WGAN-GP

Conversely, Wasserstein GANs (WGANs) [21] are an alternative solution to DCGANs, where it is introduced a clipping to the weights and the training is asynchronous (*i.e*., for each training iteration of the generator, the discriminator trains *N* iterations). Alongside, WGAN-GP [13] was proposed as an improvement of the WGAN. This updated version of the WGAN introduces a penalty term in the discriminator as an alternative to the previous weight clipping (*i.e*., a gradient penalty (GP)). This aims to “penalize the norm of the gradient of the critic’s output with respect to its input” and has shown to improve sample quality and training time [13].

### C. GAN evaluation

To use synthetic images in analysis tools and models, it is of high importance that the images are as realistic and similar to the original data as possible [10]. Nonetheless, the human evaluation of such data is time-consuming, subjective, and error-prone, which prompts the need for an evaluation metric to assess image similarity [1], [3]. Despite the increasing use of GANs, evaluating their results remains a difficult task [9]. Several measures and scores have been proposed that attempted to perform a quantitative or qualitative evaluation separately. However, to the best of our knowledge, there is no consensus on which metric is best [8]–[10].

Two of the most commonly used metrics are the Inception Score [22] and the Fréchet Inception Distance (FID) [23], which rely on the Inception V3 network [12] pre-trained on ImageNet [5] to extract the underlying characteristics of the data [4], [9], [10]. The main goal of image generation is to obtain images as similar as possible to the real data, so the original samples should be used for comparison in GAN evaluation. Nevertheless, the Inception Score does not use the original images to evaluate the synthetic ones, which is considered a limitation [23]. The FID, in turn, measures the distance between the real data distribution and the generated data distribution by calculating the mean and covariance of the activations in the final block of the InceptionV3 for both sets of images [1], [9], [23]. Therefore, lower FID scores reveal a smaller distance between the two distributions, with 0.00 being the best value (meaning that the two image sets are identical) [9], [10]. Moreover, FID is more consistent regarding image noise, artifacts, and human judgment than Inception Score and performs well as far as discriminability, computational efficiency, and robustness are concerned [9], [23]. Nonetheless, FID does not consider the fidelity and diversity of data [11], with this last one being one of the metrics used for understanding if the trained model collapsed (*i.e*., the generated images present low diversity).

## III. Methods

### A. Datasets

We used three image sets obtained from the Brain Tumor Segmentation (BraTS) 2020 and 2021 datasets [14]–[17]: Set 1 from BraTS 2020 and Sets 2 and 3 from BraTS 2021. Set 1 is the same as in our previous work [7] and presents 720 images of the FLAIR contrast. For Sets 2 and 3, we selected volumes from 48 subjects. Set 2 includes, for each subject, the slices that better represent the brain and tumor structures (central slices, #065 to #094) of the three MRI contrasts (T1, T2, and FLAIR), yielding a total of 4230 images. Finally, Set 3 is composed of slices #052 to #115 (of FLAIR contrast only) of each subject, resulting in a set of 3072 images. This set was created to broaden the cerebral area included in model training.

### B. Implementation Details

Both DCGAN and WGAN-GP were implemented using TensorFlow [24] with the Keras API [25], and their architecture follows the one implemented on [7] regarding number and topology of layers. However, we diverge from that initial architecture regarding activation functions, in which we now resort to *Leaky ReLU* with *α* = 0.2 on the Discriminator layers where in [7] *ReLU* had been applied. Regarding hyperparameters, a summary is shown in Table I. In specific, *Lr* represents the learning rate, and the optimizer was Adam [26] with *β*_1_ = 0.5 on both models. The GP term in WGAN-GP was set to 5.0, and the Discriminator trained 3 iterations for each Generator iteration.

**Table I.** Hyperparameters used in the training process.

| Model | Image Set | Epochs | Lr | Batch size | Dropout |
| --- | --- | --- | --- | --- | --- |
| DCGAN | Set 1 | 50 | 0.0001 | 8 | 0.2 |
|  | Set 2 | 100 | 0.0002 | 64 | 0.5 |
|  | Set 3 | 100 | 0.0002 | 32 | 0.5 |
| WGAN-GP | Set 1 | 50 | 0.0005 | 8 | 0.2 |

The DCGAN model was trained with the three image sets, while the WGAN-GP was only trained with image Set 1 since it could not handle training with larger image sets.

### C. Setup

The models were trained in 3 nodes (of a 4-node cluster with the Simple Linux Utility for Resource Management (SLURM) job management system [27]), each containing one NVIDIA GeForce RTX 2080 Ti GPU. The distribution strategy was TensorFlow’s `MultiWorkerMirroredStrategy`, which implements synchronous training where the steps are synced across the workers and replicas. As such, all workers train over different slices of input data and aggregate gradients at each step.

## IV. Results and Discussion

Our models generate one 256*×*256 brain MRI image (axial view) per epoch of training. We measured the FID score for each generated image set with InceptionV3 [12] pre-trained on ImageNet [5] and with RadImageNet [6] weights. The top 3 FID values obtained for each model are presented in Table II, where FID_ImageNet_ refers to FID calculated with pre-training on ImageNet and FID_RadImageNet_ means FID calculated with RadImageNet weights.

**Table II.**
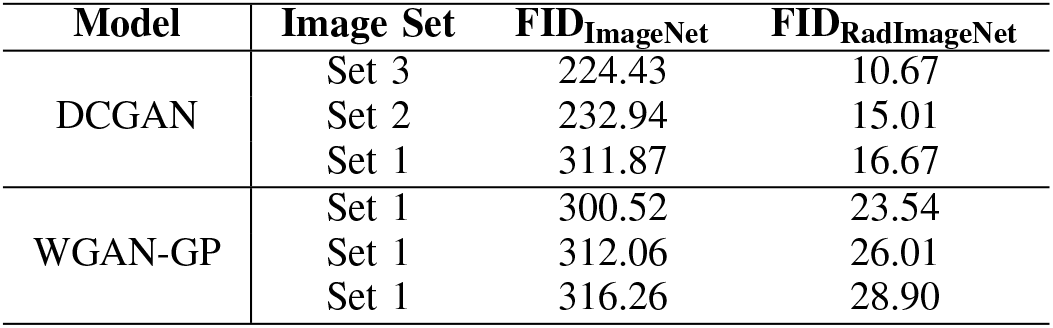
Top 3 FID scores for the two models.

In general, DCGAN FID values are lower than those of WGAN-GP, and evaluating with ImageNet weights scores higher FID than with RadImageNet. This goes in line with the literature [3], [4] and can be explained by the difference in the pre-training datasets: ImageNet is a non-medical dataset, while RadImageNet contains labeled images from several imaging modalities (PET, CT, Ultrasound, and MRI) [6], so applying RadImageNet weights allows the InceptionV3 to grasp features specific to medical images. The FID_ImageNet_ values obtained with DCGAN, although significantly high, fall into what other author’s findings show (*i.e*., FID roughly between 50 and 270) [1], [2], so we theorize that our FID_ImageNet_ WGAN-GP values are also coherent considering the use of the ImageNet dataset in the medical context. Furthermore, the FID_RadImageNet_ scores of both models are consistent with results from other works [4], [8], with the best values ranging from 10.67 to 28.90, representing a decrease of up to 21*×* when compared to the FID results obtained with ImageNet.

As reported in [1], we found that the larger the image set given to the GAN, the better the FID score of the synthetic images will be (Set 3 has more images than Sets 1 and 2 and gave the best FID score) and, consequently, the better the model will perform. Moreover, we report better FID values with DCGAN trained on BraTS 2021 image sets (Sets 2 and 3) than those obtained in [18] with other GAN architectures. All these findings suggest that our models have a similar performance in a distributed environment to those of others in a non-distributed setting, which supports our goal of applying GANs in an HPC environment to generate training data for other ML/DL models. The best results obtained for Sets 1, 2, and 3 are shown in Figures 1, 2, and 3, respectively.

**Fig. 1.**
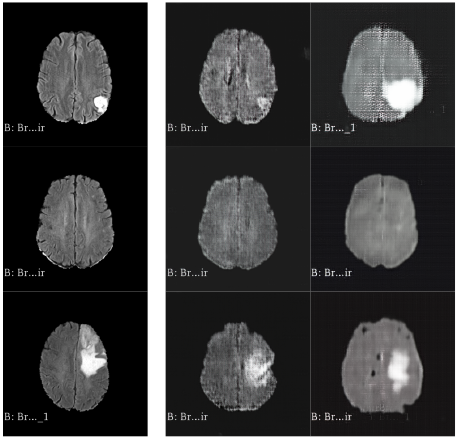
Best results obtained for Set 1 (by columns, from left to right: original images, DCGAN, WGAN-GP).

**Fig. 2.**
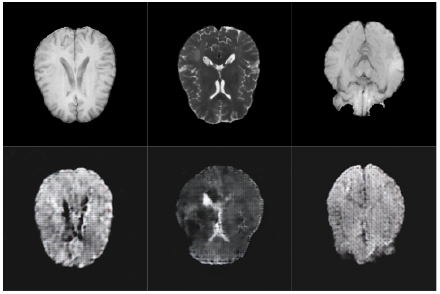
Best results obtained for Set 2 (upper row contains original images, lower row contains DCGAN images).

**Fig. 3.**
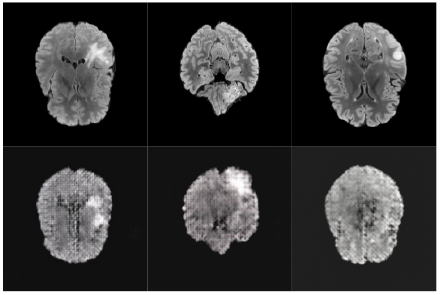
Best results obtained for Set 3 (upper row contains original images, lower row contains DCGAN images).

As seen in Figures 1-3, the generated images have, overall,less detail than the original samples. Conversely, despite our models’ performance in all contrasts, two out of three of our image sets were from FLAIR contrast, so more tests with other contrasts are needed to assess the model’s generalization ability. Finally, the WGAN-GP raised issues regarding the training with Sets 2 and 3, which may occur because this model has a slower convergence than DCGAN [13] and training with a larger image set implies learning a higher amount of characteristics. Nevertheless, more tests are needed to pinpoint the exact reason for the WGAN-GP training problems.

## V. Conclusions and Future Work

Image generation has turned into a powerful mechanism to overcome the scarcity of medical data for training, with GANs leading in terms of synthetic image quality. Since training these models is computationally demanding and requires a large amount of memory [1], workload distribution becomes the next big step for medical image generation.

In this work, we re-implemented two GANs (a DCGAN and a WGAN-GP) in a distributed setting to generate MRI images of the brain. To evaluate the synthetic images, there is no consensus on which metric is best; each one captures different aspects of the image generation process, so it becomes unlikely that a single measure can encompass all aspects [8]–[10]. We evaluated our results using FID with InceptionV3 pre-trained on ImageNet and RadImageNet weights.

We obtained FID scores consistent with the literature, either with ImageNet or RadImageNet weights, and FID_RadImageNet_ was significantly lower than FID_ImageNet_. Moreover, we validated that a larger input image set results in better model performance, as the FID scores decreased with Sets 2 and 3. With this, we present evidence that it is possible to obtain similar image quality and model performance using distributed environments, and although we recognize that applying FID to GAN evaluation in medical imaging raises some questions (concerning InceptionV3 being trained with representations of a non-medical dataset) [1], [3], [4], [9], we believe that FID with RadImageNet weights is a strong metric of image quality, as our findings match those of other works in the same scope. Future work on this approach includes testing the models in a larger distributed setting, more experiments to fine-tune the hyperparameters (*i.e*., to improve synthetic image quality), and further investigation on the WGAN-GP training process.

